# Gamma-protocadherin localization at the synapse corresponds to parameters of synaptic maturation

**DOI:** 10.1101/760041

**Authors:** Nicole LaMassa, Hanna Sverdlov, Aliya Mambetalieva, Stacy Shapiro, Michael Bucaro, Monica Fernandez-Monreal, Greg R. Phillips

## Abstract

Clustered protocadherins (Pcdhs) are a large family of ~60 cadherin-like proteins (divided into the subclasses α, β, and γ) that compose a surface “barcode” in individual neurons. The code is generated through combinatorial expression via epigenetic regulation at a large gene cluster that encodes the molecules. During early neural development, Pcdhs were shown to mediate dendrite self-avoidance in some neuronal types through a still uncharacterized anti-adhesive mechanism. Pcdhs were also shown to be important for dendritic complexity in cortical neurons likely through a pro-adhesive mechanism. Pcdhs have also been postulated to participate in synaptogenesis and the specificity of connectivity. Some synaptic defects were noted in knockout animals, including synaptic number and physiology, but the role of these molecules in synaptic development is not understood. The effects of Pcdh knockout on dendritic patterning may present a confound to studying synaptogenesis. We have shown previously in vivo and in cultures that Pcdh-*γ*s are highly enriched in intracellular compartments located in dendrites and spines with localization at only a few synaptic clefts. To gain insight into how Pcdh-*γ*s might affect synapses, we compared synapses that harbored endogenous Pcdh-*γ*s versus those that did not for parameters of synaptic maturation including pre- and postsynaptic size, postsynaptic perforations, and spine morphology by light microscopy in cultured hippocampal neurons and by serial section immuno-electron microscopy in hippocampal CA1. In mature neurons, synapses immunopositive for Pcdh-*γ*s were found to be larger in diameter with more frequent perforations. Analysis of spines in cultured neurons revealed that mushroom spines were more frequently immunopositive for Pcdh-*γ*s at their tips than thin spines. Taken together, these results suggest that Pcdh-*γ* function at the synapse may be related to promotion of synaptic maturation and stabilization.

## Introduction

The specificity of wiring of the vertebrate nervous system is achieved at several levels or stages during embryonic and postnatal development. Postmitotic neurons must elaborate dendritic arbors in the correct spatial orientation as well as send axons out along proper trajectories. Later in development, axons interact with the dendrite to form excitatory synapses. In large part, these developmental processes are facilitated by the action of cell-surface adhesion and recognition proteins (Kim, 2019) that allow neuronal outgrowths to sense their environment and respond accordingly.

It has been proposed that cell adhesion molecules also establish a system of suitable specificity that can restrict interactions between the enormous number of neurons and glial cells. One group of cell adhesion molecules, the clustered protocadherins (Pcdhs), a family of ~60 adhesion molecules encoded on a large gene cluster (Wu and Maniatis, 1999; Zipursky and Sanes, 2010), fulfills these criteria. Members of the three subclasses of Pcdhs (α, β, and γ) are expressed stochastically through epigenetic regulation (Guo et al., 2012; Hirayama et al., 2012; Monahan et al., 2012; Jiang et al., 2017). A combination of molecules from the three subclasses generates a unique homophilic recognition unit (Schreiner and Weiner, 2010; Goodman et al., 2016, 2017; Brasch et al., 2019) on neuronal surfaces.

In vertebrates, these cell-specific recognition units, presumed to be identical on the surfaces of same-cell dendrites, most likely are primary mediators of dendritic self-avoidance in certain neurons that have planar dendrites such as the retinal starburst amacrine cell and the cerebellar Purkinjie cell, as demonstrated in Pcdh-*γ* conditional null animals (Lefebvre et al., 2012; Ing-Esteves et al., 2018). Other roles for Pcdhs in dendrite elaboration (Suo et al., 2012) have been observed. This role does not appear to involve same-cell recognition but rather astrocyte-dendrite interaction (Garrett and Weiner, 2009; Molumby et al., 2016) in cortical neurons. It is possible that there are different modes of Pcdh interactions, those between same cell processes and those between different cells and that these modes may have different outcomes in terms of positive or negative effects on cell adhesion.

Overall, the roles of Pcdhs in early neurodevelopmental events, such as dendrite (Garrett and Weiner, 2009; Molumby et al., 2016) and axon (Hasegawa et al., 2008, 2012, 2016; Prasad and Weiner, 2011; Chen et al., 2017; Katori et al., 2017; Lu et al., 2018) outgrowth, have been the most studied. However, Pcdhs remain expressed long after these events have ended (Phillips et al., 2003; Frank et al., 2005). Pcdhs are located at some excitatory synapses at 1 month postnatally in rodents at the synaptic cleft and within intracellular synaptic compartments in vivo and in cultures (Wang et al., 2002; Phillips et al., 2003; Fernández-Monreal et al., 2009, 2010). Pcdhs have been shown to participate in synaptic development but the exact role may depend on cell type. For example, mice lacking Pcdh-*γ*s exhibit deficits in synaptic numbers and physiology in spinal cord neurons (Weiner et al., 2005) but increased synapses in cortical neurons with an increase in the number of thin, immature spines (Molumby et al., 2017). In contrast, other groups reported that Pcdh-α and -γ knockout or knockdown caused reduced synaptic numbers in hippocampus (Suo et al., 2012). Knockout of a subset of Pcdh-*γ*s expressed constitutively in all neurons (Pcdh-γC3-5) reduced the density of inhibitory synapses in cultured hippocampal neurons (Li et al., 2012). A possible confound to studying synaptogenesis in these knockout experiments is that the reduced dendritic complexity due to Pcdh knockout might itself affect synaptic density, thus necessitating a different approach to confirm a role for Pcdhs at the synapse. Pcdh-*γ*s can interact with various synaptic and signaling molecules (Chen et al., 2009; Han et al., 2010; Garrett et al., 2012; Keeler et al., 2015; Mah et al., 2016; Molumby et al., 2017; Fan et al., 2018) but which of these coordinate with Pcdhs to affect synaptic development is still unknown. It has been hypothesized that Pcdhs play a role in late synaptic developmental events (Jontes and Phillips, 2006).

It is known that synaptic stability and maturation are associated with a widening of the synapse and dendritic spine head (reviewed in (Berry and Nedivi, 2017)). The appearance of perforations in the postsynaptic density has also been associated with an increased number of AMPA receptors which is suggestive of a mature and strengthened synapse (Desmond and Weinberg, 1998; Lüscher et al., 2000; Nicholson et al., 2006). Synaptic perforations have also been associated with LTP induction (Toni et al., 2001).

Antibodies that detect Pcdh-*γ*s label striking punctate structures in vivo and in cultured hippocampal neurons that are thought to represent trafficking organelles but also, at times, overlapped with synaptic markers (Wang et al., 2002; Phillips et al., 2003; Fernández-Monreal et al., 2009, 2010). We asked whether synapses positive for Pcdh-*γ*s exhibit differences in terms of diameter and perforation status relative to unlabeled synapses. Using serial section immuno-electron microscopy in vivo and confocal microscopy of immunolabeled cultured neurons we compared synapses that harbor Pcdh-*γ*s to those that do not for these parameters of synaptic maturation. In mature neurons, we found that Pcdh-γ positive synapses were significantly larger and were more likely to be perforated. When Pcdh-*γ*s were located near the tip of the dendritic spine, that spine was more likely to be of the mushroom type. These results suggest that Pcdh-*γ*s could be associated with stabilization and maturation events of excitatory synapses in hippocampus.

## Materials and Methods

### Hippocampal neuron cultures and labeling

Neurons were cultured from embryonic day 18 rat embryos as described previously (Phillips et al., 2003; Fernández-Monreal et al., 2009) and grown for 12 or 21 days. In some cases, neurons were transfected at day 15 with pEGFP-n1 to fill dendrites with GFP and grown for an additional 7 days as described (Fernández-Monreal et al., 2009). For synaptic measurements, neurons were fixed and immunolabeled with an affinity purified antibody raised against the Pcdh-*γ* constant cytoplasmic domain (Phillips et al., 2003). This antibody was validated in (Fernández-Monreal et al., 2010) by showing that it labels the same puncta as labeled with a commercially available monoclonal antibody to the Pcdh-*γ* constant domain (clone 195/5, UC Davis/NIH NeuroMab Facility). For synaptic labeling, neurons were colabeled with goat anti-PSD-95 (Abcam Cat# ab12093, RRID:AB_298846) and mouse anti-vGlut (Millipore Cat# MAB5502, RRID:AB_262185). For spine measurements, GFP filled neurons were stained with anti-Pcdh-γonly.

### Confocal image analysis

Images were obtained with a Zeiss LSM510 META confocal microscope. Double and triple label z-stack images were acquired in multi-track mode at 4 times line averaging with a 63x/1.4 NA oil DIC objective. Z-stacks were flattened by maximum projection for image analysis. Images were imported into NIH Image J as separate channels for colocalization analysis. 100 distinct synapses were mapped and assigned a region of interest (ROI) along the length of the dendritic tree in each neuron using the anti-vGlut/PSD-95 colocalized images. Then, the Pcdh-*γ* channel was merged separately with the PSD-95 channel or the vGlut channel. Pixels in which Pcdh-*γ* colocalized with either synaptic marker were detected with the RG2B colocalization plugin which generates a new image of the colocalized pixels. The colocalization images were used to map and save regions of interest (ROIs) using the ROI manager in Image J. Synapses that did not show Pcdh-γ colocalization with PSD-95 and/or vGlut were also mapped. To identify synapses in which Pcdh-*γ*s overlapped simultaneously with PSD-95 and vGlut, images representing the colocalized pixels from Pcdh-γ/PSD-95 and Pcdh-γ/vGlut were loaded into separate channels and subjected again to the RG2B colocalization plugin which resulted in an image map containing pixels of colocalization for all three markers. ROIs of synapses from Pcdh-γ/PSD-95, Pcdh-γ/vGlut and Pcdh-γ/PSD-95/vGlut colocalization, as well as unlabeled synapses, were saved and remapped onto the original single channel PSD-95 or vGlut images. The area and average pixel value of the synaptic PSD-95 and vGlut punctum was determined in Image J for all synapses and averaged over 3 neurons.

### Animals

All animals used in this study were in accordance with the Institutional Care and Use Committee (IACUC). Postnatal day 30 Sprague-Dawley rats were prepared for electron microscopy by freeze substitution and low temperature embedding as described previously (Fernández-Monreal et al., 2010).

### Serial sectioning

The block was roughly trimmed using a single edge, heavy duty razor blade. Then, it was finely trimmed to include a narrow area of CA1 including the pyramidal layer and stratum radiatum using a Diatome trimtool 20 (Diatome). Serial 70 nm sections were cut with a 45° diamond knife (Diatome) using a Leica UCT Ultramicrotome. Ribbons of ~5 sections were oriented and collected on formvar/carbon coated nickel slot grids (Electron Microscopy Sciences) using a perfect loop (Electron Microscopy Sciences).

### Postembed immunogold labeling and staining

Grids were loaded into a Chien staining pad (Ted Pella) and sections were immunolabeled with an affinity-purified rabbit polyclonal antibody generated against the mouse Pcdh-*γ* constant cytoplasmic domain (Phillips et al., 2003) diluted 1:5 in 20 mM Tris pH 7.4, 20 mM NaCl, 3% BSA and 1% Triton-X 100 overnight in a humidified chamber. The sections were then washed three times with 20 mM Tris pH 7.4, 20 mM NaCl and incubated with secondary 15 nm gold-tagged goat-anti-rabbit IgG F(ab’)2 (Electron Microscopy Sciences) diluted 1:10 in 20 mM Tris pH 7.4, 20 mM NaCl, 3% BSA and 1% Triton-X 100, overnight in a humidified chamber. The grids were washed again three times and then rinsed once with deionized water.

The sections were stained with 4% uranyl acetate for 40 minutes in a dark chamber and washed three times with deionized water. The grids were then stained with Reynold’s lead citrate for 30 seconds and washed three times with deionized water. The grids were left to dry before imaging.

### Acquisition of serial images and EM image analysis

Serial images of CA1 stratum radiatum were acquired using a Fei Tecnai Spirit Transmission Electron Microscope (Advanced Imaging Facility, College of Staten Island) at 18,500x magnification. Identification of corresponding areas of stratum radiatum in each consecutive section was facilitated using blood vessels as landmarks.

Image series were saved as stacks in Image J. Every synapse in each stack was identified, the diameter of the postsynaptic density was measured using the section through the widest part of the synapse in the serial images. Synapse status as macular or perforated was compiled visually from the serial images. A synapse was considered labeled if it contained at least two gold particles on its cleft, spine or axonal compartment in a section. The average diameter of labeled or unlabeled synapses was compared and statistical significance determined by t test.

### Spine analysis

Maximum projection confocal images of 3 GFP transfected and anti-Pcdh-*γ* labeled neurons were imported into NIH image J. Only isolated spines were analyzed for Pcdh-*γ* labeling. Pcdh-*γ* positive and negative spines were classified as either thin, stubby, or mushroom shaped using criteria described previously (Harris et al., 1992). Pcdh localization was investigated by the presence of pixels located at the head, neck, or base of the spine. Diffuse labeling was not considered. Pcdh localization in relation to spine shape was assessed and was compared to unlabeled spines. t tests were conducted for significance.

## Results

### Pcdh-*γ* localization at synapses is associated with larger synaptic puncta

Previous studies have shown Pcdh-*γ*s to be localized at a subset of synapses in vivo (Wang et al., 2002) and in cultured neurons (Phillips et al., 2003) but the role of these molecules in synaptic development remains unknown. Some defects in synaptic number and physiology in in Pcdh-*γ* null animals have been noted (Weiner et al., 2005; Suo et al., 2012; Molumby et al., 2016) suggesting the possibility that Pcdh-*γ*s might participate in synaptic maturation or stability. Synapse size is one parameter that can be associated with synaptic stability (Berry and Nedivi, 2017). To compare synapses that harbor Pcdh-*γ*s to those that do not, we immunostained cultured hippocampal neurons at 2 developmental stages (days 12 and 21 in vitro) with anti-Pcdh-γ antibodies together with antibodies to PSD-95 and vesicular glutamate transporters (vGlut) (Fig. 1a). From day 12 to day 21 there was a dramatic increase in the average number of synapses per neuron indicating ongoing synaptogenesis during this time period (Fig. 1b).

**Figure 1.**
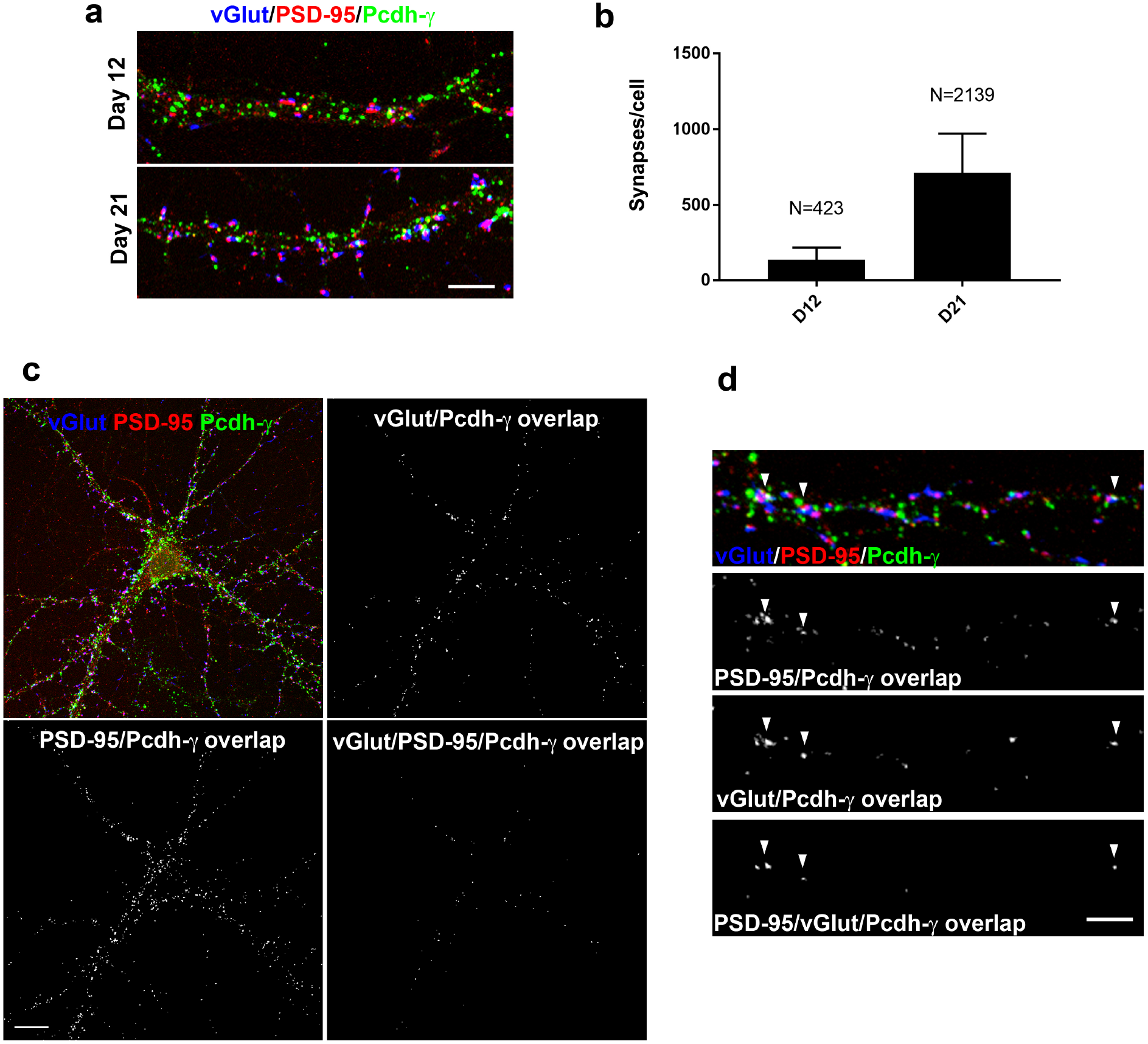
Unbiased selection of Pcdh-*γ* positive synapses in hippocampal neurons triple-labeled with antibodies to vesicular glutamate transporter (vGlut; blue channel), PSD-95 (red channel), and anti-Pcdh-*γ* constant domain (green channel). (a) Dendrite segments from neurons at day 12 and day 21. Synapses occasionally overlapped with Pcdh-γpuncta at both stages. (b) The numbers of synapses per neuron increased from day 12 to day 21 indicating ongoing synaptogenesis. (c) Low magnification image of a single labeled neuron at day 21 and selection for Pcdh-*γ* positive synapses (top left). Image generated in which Pcdh-*γ* signal overlaps with that of vGlut (top right). Image generated in which Pcdh-*γ* signal overlaps with PSD-95 (bottom left). vGlut/Pcdh-*γ* and PSD-95/Pcdh-*γ* overlapping pixels were loaded into the red and green channels of a new image and overlapping pixels selected (bottom right). (d) Higher magnification of a dendrite and selection of pixels corresponding to Pcdh-*γ* label that overlaps with vGlut, PSD-95, or both simultaneously. Arrowheads indicate synapses with simultaneous Pcdh-*γ* colocalization with pre- and postsynaptic markers. Bar = 5 μm in (a) and (d) and 10 μm in (c).

We randomly sampled ~100 synapses from 3 neurons each at days 12 and 21 and used an unbiased selection method based on overlap of signal from Pcdh-*γ* immunolabeling with the signal for both pre and postsynaptic makers to identify synapses with overlapping Pcdh-γsignal in day 12 and day 21 cultured hippocampal neurons (Fig. 1c and d). The selection method identified synapses that exhibited Pcdh-*γ* puncta overlapping with a presynaptic or postsynaptic punctum individually or simultaneously.

We compared pre- and postsynaptic punctum area for three categories of synapses (Fig. 2a). Synapses with no overlapping Pcdh-*γ* immunoreactivitiy were considered “unlabeled”. Synapses with an overlapping Pcdh-*γ* punctum either on the presynaptic or postsynaptic specialization, or different Pcdh-*γ* puncta overlapping with the presynaptic and postsynaptic specializations at a single synapse, but never simultaneously overlapping with both, were considered labeled in an “uncorrelated” fashion. Finally, synapses with at least one single Pcdh-*γ* punctum simultaneously overlapping with both pre- and postsynaptic specializations were considered labeled in a “correlated” manner (Fig. 2a). Thus, “correlated” labeling may be considered to be the most stringent filter for identification of Pcdh-*γ* positive synapses, and this labeling may correspond to synaptic compartment or cleft labeling.

**Figure 2.**
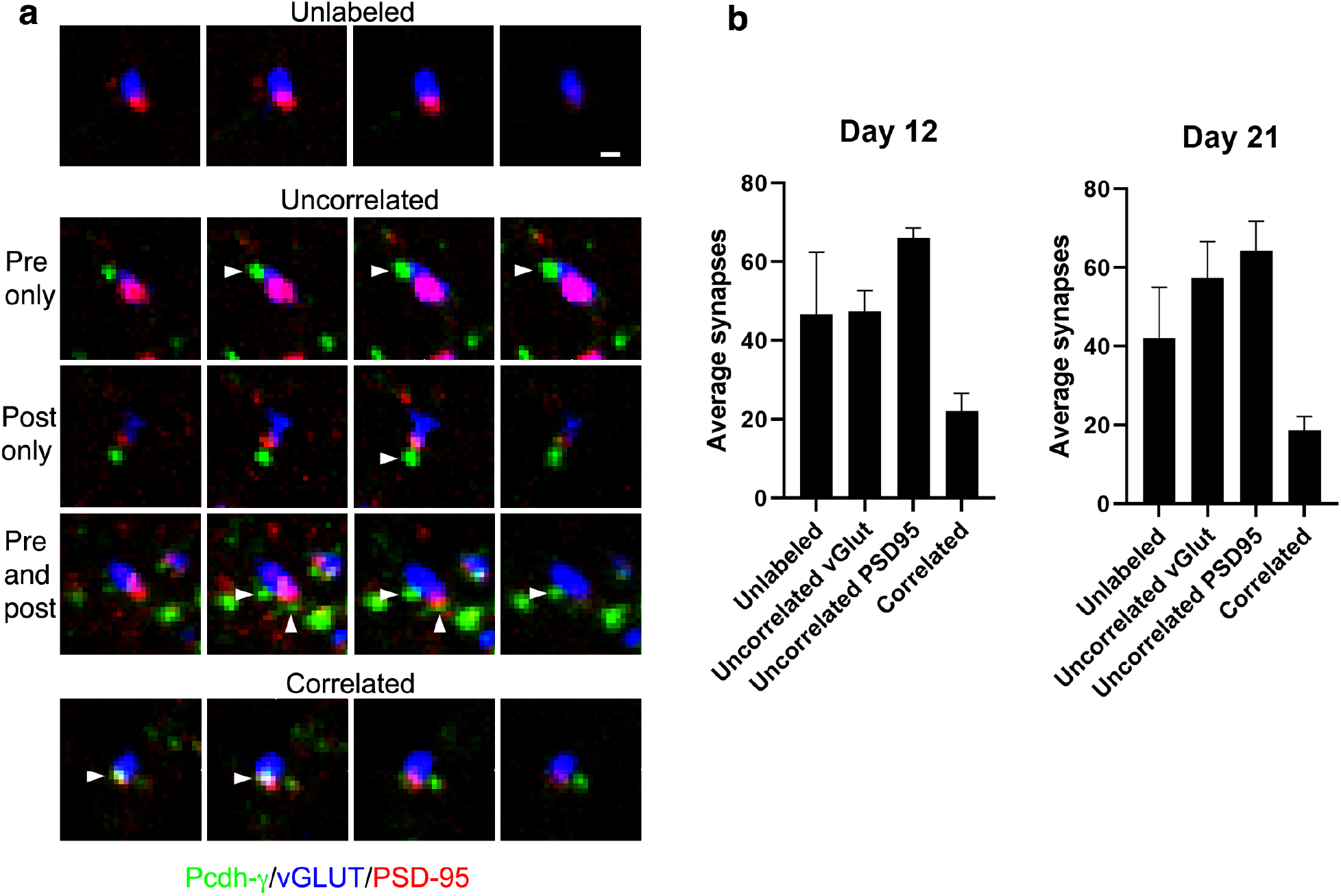
Pre- and postsynaptic profiles for Pcdh-*γ* negative synapses or synapses in which Pcdh-*γ* label overlaps with one or both synaptic markers simultaneously. (a) Examples of synapses from day 21 cultures in which Pcdh-*γ* signal does not overlap with either synaptic marker (unlabeled), or where Pcdh-*γ* signal overlaps with one or both synaptic markers (middle, arrowheads) but not simultaneously (uncorrelated), or synapses with a Pcdh-*γ* punctum overlapping (bottom, arrowhead) with both synaptic markers simultaneously (correlated). (b) Quantification indicates similar proportion of synaptic labeling at day 12 and day 21. Many uncorrelated synapses were counted twice due to the fact that Pcdh-*γ*s can overlap with one or both synaptic markers at the same synapse.

The percentage of synapses with Pcdh-*γ*s colocalizing with one or both synaptic markers was similar at both stages (Fig. 2b). These percentages added up to > 100% due to the fact that Pcdh-*γ* puncta could be colocalized with both pre- and postsynaptic markers either separately or simultaneously (see Fig. 2).

The area of synaptic puncta was determined in Image J from un-resampled images (Fig. 3a). At day 12, there was no change in the average area or pixel intensities for both presynaptic and postsynaptic puncta over the three categories of synapses (Fig. 3b, top). However, at day 21, at synapses with a correlated Pcdh-*γ* label, there was a significant increase in the area of both the pre- and postsynaptic puncta without a change in the average pixel values (Fig. 3b, bottom). The specific effect at day 21 suggests Pcdh-*γ*s, when present either at the synaptic cleft or within synaptic compartments near the cleft, as has been reported previously (Fernández-Monreal et al., 2010), might be indicative of a synaptic maturation process late in synaptogenesis.

**Figure 3.**
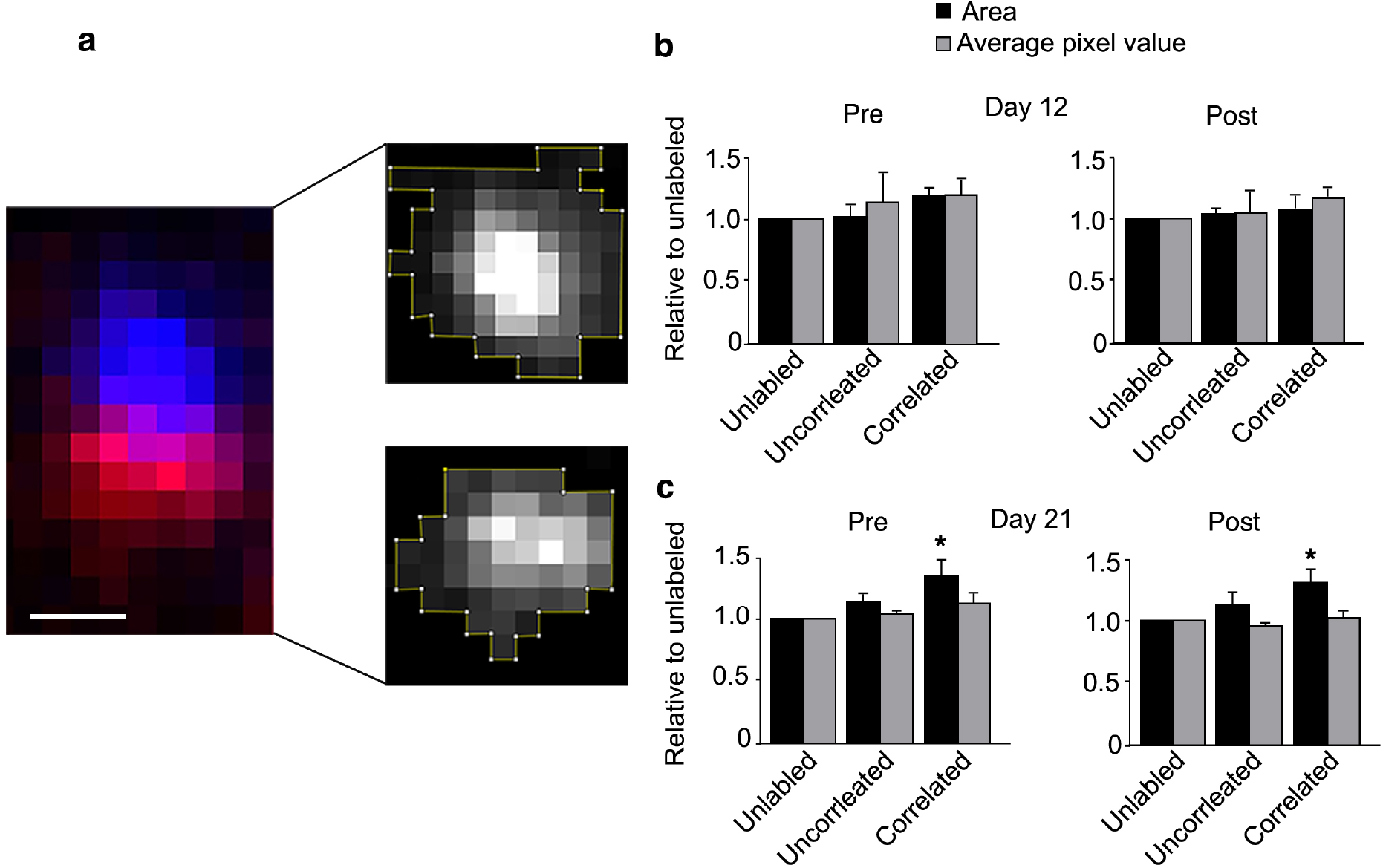
Quantitative analysis of the area and the average pixel value of Pcdh-*γ* positive and negative synapses. (a) A synapse from a confocal projection immunolabeled with antibodies to vGlut (blue pixels), PSD95 (red pixels). Channels were separated for quantification of area and average pixel intensity in Image J (white pixels). (b) Quantification of area and average pixel values for pre-(left) and postsynaptic (right) puncta for the three categories of synapses at day 12 and 21. There was no difference in area and pixel values for both pre- and postsynaptic markers for the 3 categories of synapses at day 12. However, at day 21, when Pcdh-*γ* correlates with both pre- and postsynaptic puncta simultaneously there was a significant increase in the area of both puncta (p<0.05). Bar = 300 nm.

### Serial immuno-EM analysis of anti-Pcdh-*γ* labeled synapses in vivo

The light level observations suggested that synaptic Pcdh-*γ*s delinate maturing synapses with larger synaptic specializations. To study this in more detail in vivo, Pcdh-*γ* antibodies were used to label serial sections of rat hippocampal CA1 at 30 days postnatal, which is a period of ongoing synaptic development (Harris et al., 1992). Using serial sectioning, labeled synapses can be identified and synaptic parameters, such as the diameter of the synapse, as well as perforation status, accurately quantified by scanning through the entire width of the synapse. In many instances, gold particles labeled the same structure in multiple sections (arrows and double arrowheads, Fig. 4). Overall, Pcdh-*γ* labeling was found at a few synaptic clefts (double arrowheads, Fig. 4), synaptic (double arrows, Fig. 4) and non-synaptic (arrowheads, Fig. 4) intracellular compartments and at non-synaptic cell-cell contact points (asterisk, Fig. 4), as previously reported (Phillips et al., 2003; Fernández-Monreal et al., 2010).

**Figure 4.**
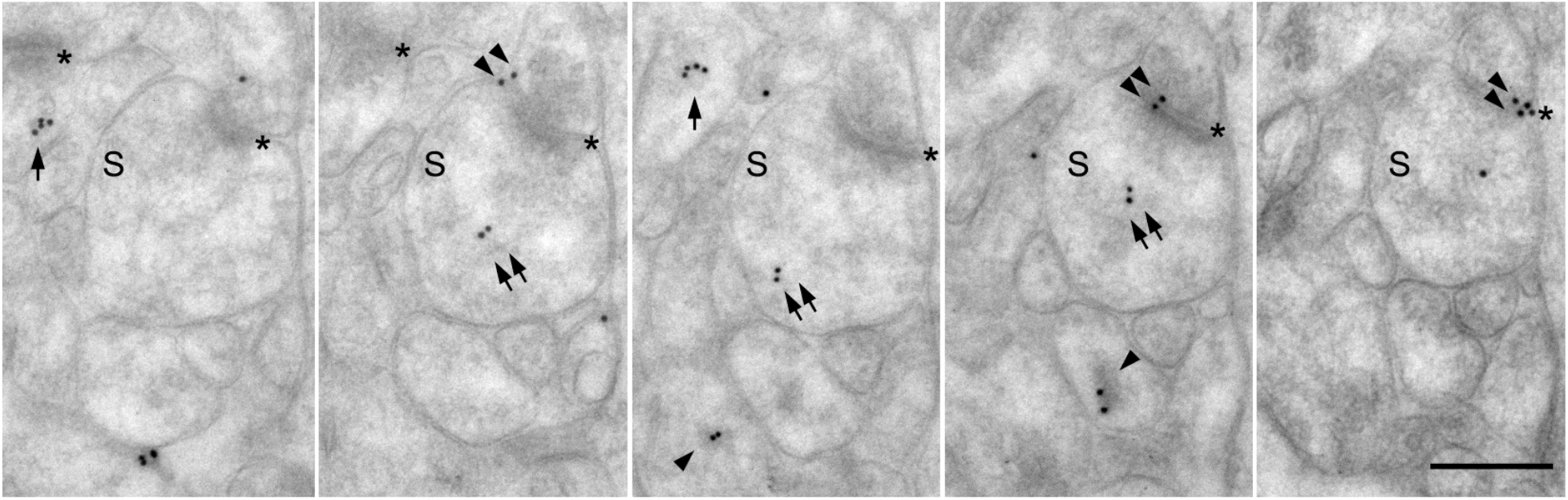
Immuno-EM serial section analysis of synapses labeled with anti-Pcdh-γ. Five serial sections through a synapse positive for Pcdh-*γ*s at the cleft (double arrowheads) and an internal dendritic compartment (double arrows). Single arrows show a synaptic compartment labeled in 2 sections. Single arrowhead shows labeled nonsynaptic compartments. Asterisk delineates synaptic clefts. Bar = 500 nm.

Quantitative comparison of anti-Pcdh-γ-labeled and unlabeled synapses was performed. The diameter of every synapse was measured through the widest part of each synapse in serial sections (n=140 synapses in 21 serial image stacks). Each synapse was also classified as either macular or perforated. Synapses were then categorized as anti-Pcdh-*γ* labeled or unlabeled. The majority of Pcdh-*γ* labeled synapses were immunopositive for Pcdh-*γ* within intracellular compartments as previously described (Fernández-Monreal et al., 2010). Therefore synapses were considered as labeled if multiple gold particles were present over the same subcellular structure either at the cleft or within an dendritic or axon compartment, or some combination thereof (Fig. 5a). Overall, it was found that Pcdh-*γ* labeled synapses exhibited an average diameter of 484 nm which was significantly larger when compared to unlabeled synapses, that averaged 222 nm in diameter (Fig. 5b).

**Figure 5.**
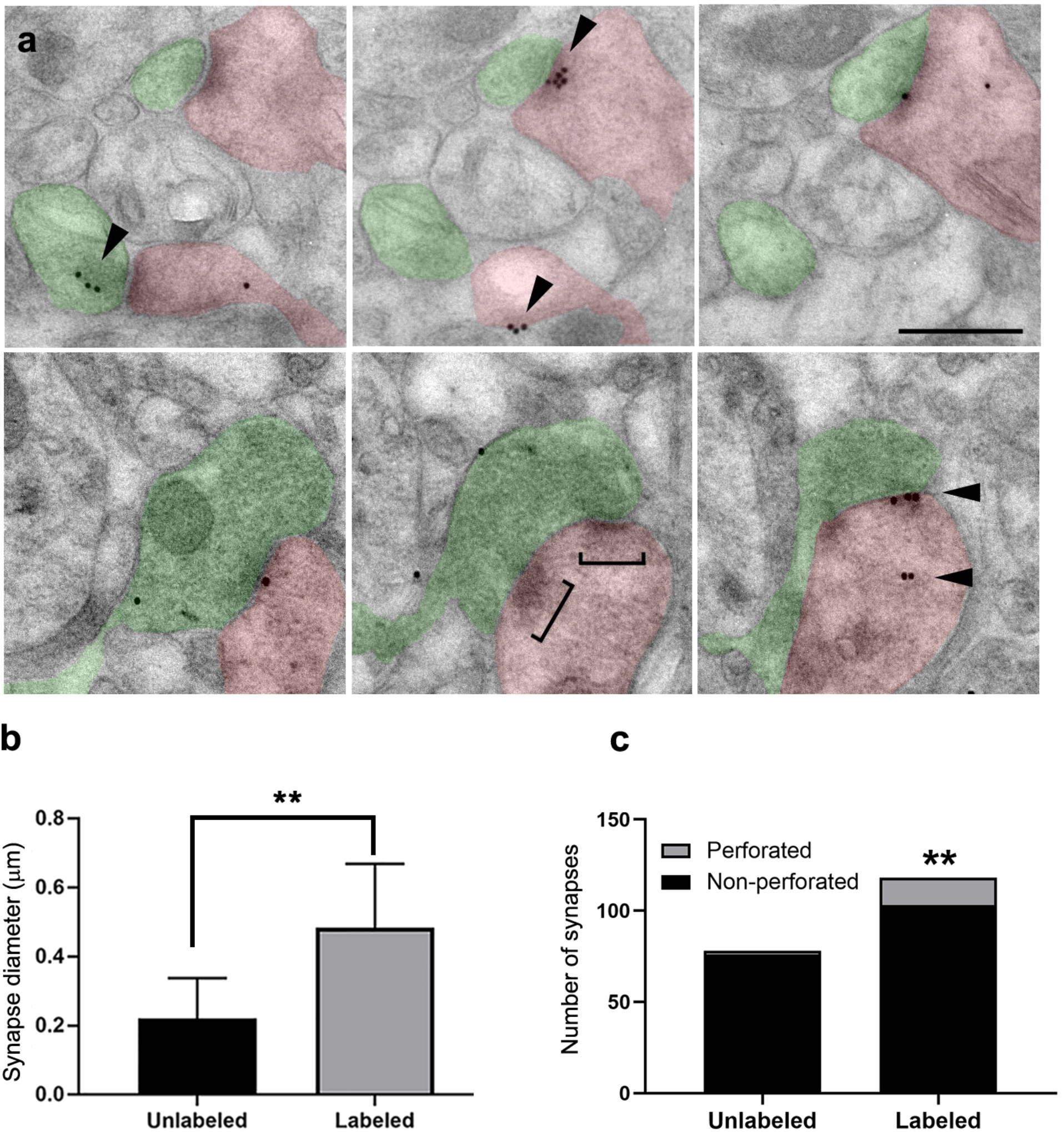
Pcdh-*γ* synapses have larger diameter and are more likely to be perforated. (a) Serial sections through Pcdh-γ positive synapses. Pcdh-*γ* labeling at clefts and synaptic compartments indicated by an arrowhead. Some Pcdh-*γ* positive synapses were macular (top) while others were revealed to be perforated (brackets, bottom) upon serial sectioning. Bar = 500 nm. (b) Quantification of synapse diameter for Pcdh-*γ* labeled versus unlabeled synapses. Unlabeled synapses averaged 200 nm in diameter while labeled synapses were more than twice as large at 400 nm in diameter (p<0.005). (c) The overwhelming majority of unlabeled synapses were macular type synapses while ~20% of labeled synapses were perforated (chi-squared test=0.01).

To determine if Pcdh-*γ* positive synapses are more likely to be perforated, synapses were classified as macular or perforated through examination of serial sections (Fig. 5a). Of the 78 unlabeled synapses, only 2 were perforated, while in contrast, among the 106 labeled synapses, 16 were perforated, which was a signficant difference (Fig. 5c).

### Pcdh-*γ* position in dendritic spine heads corresponds to a mushroom type

The Pcdh-*γ* immunostaining pattern at the light level is strikingly discrete, with puncta that can localize at the spine head, neck or dendritic shaft (Fernández-Monreal et al., 2009). Mushroom type spines are thought to be the most stable spines while thin spines are more transient (reviewed in (Berry and Nedivi, 2017). To examine the relationship between Pcdh-*γ* expression at the spine and spine morphology, we transfected primary neurons with GFP which were then grown for 21 days, fixed and immunostained with anti-Pcdh-γ (Fig. 6a). All spines were first categorized as being either mushroom, thin, or stubby, then the location of Pcdh-*γ* puncta was assigned to either the spine head (‘a’, Fig. 6b), the spine neck (‘b’, Fig. 6b) or the dendritic shaft at the spine base (‘c’, Fig. 6b) There were relatively few stubby spines in these neurons and these spines were therefore excluded from further analysis.

**Figure 6.**
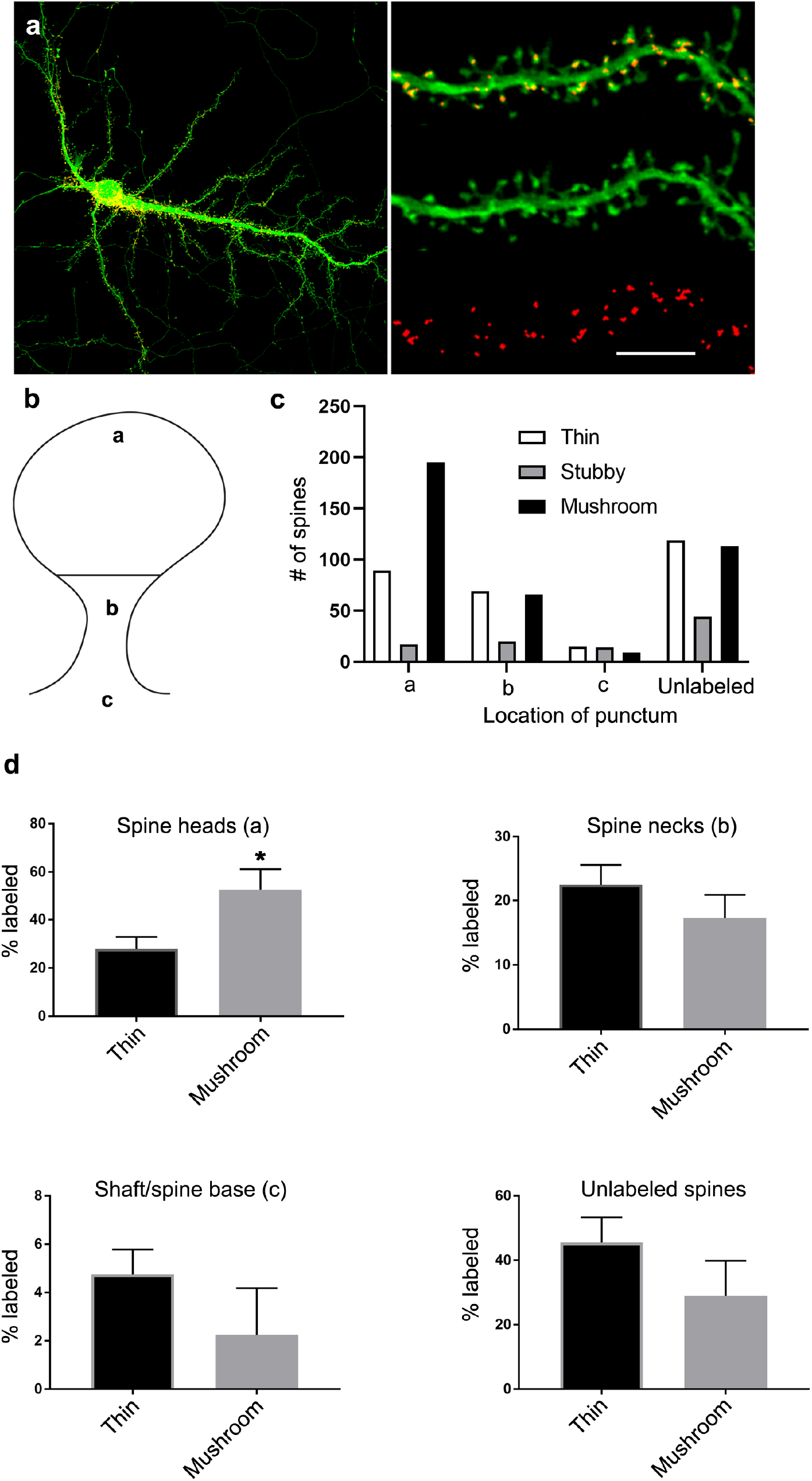
Mushroom spine heads are more likely to be Pcdh-*γ* positive than thin spine heads. (a) GFP (green channel) filled neuron labeled with anti-Pcdh-*γ* (red channel). (b) Categorization of labelling patterns based on location of Pcdh-*γ* positive puncta with the spine (‘a’ denotes spine head labeling, ‘b’, spine neck labeling and ‘c’, shaft labeling directly below the spine). (c) Distribution of spine type with respect to labeling pattern. (d) Quantification of the proportion of thin or mushroom spines that exhibit the indicated labeling pattern. The proportion of unlabeled, spine neck labeled, or dendritic shaft labeled spines was not significantly different between thin and mushroom spines while spines that were labeled on the head were more likely to be mushroom type spines (p<0.05) Bar= 10 μm.

Many spines were unlabeled, consistent with the variability of Pcdh-*γ* localization at synapses reported previously (Wang et al., 2002; Phillips et al., 2003). When present at spines, the most frequent location for Pcdh-*γ* puncta in both mushroom and thin spines was in the spine head (Fig. 6c). However, some spines were also labeled at the spine neck with a smaller number labeled at the base of the spine in the dendritic shaft. A significant proportion of spines were unlabeled (Fig. 6c).

Most (51%) of the mushroom spines were labeled at the spine head (Fig. 6d, upper left) which differed with thin spines (Fig. 6d, upper right) that were less frequently labeled at the spine head. Pcdh-*γ* labeling at spine necks or the base was not predictive of spine type; neither were unlabeled spines predominately of any one type (Fig. 6d). These results suggest that Pcdh-γfunction in the spine compartment could be associated with some aspect of spine maturation.

## Discussion

The Pcdh gene cluster is capable of generating, through epigenetic regulation of ~60 different isoforms (Guo et al., 2012; Hirayama et al., 2012; Monahan et al., 2012; Yin et al., 2017), what has been considered an adhesive “barcode” (Rubinstein et al., 2017; Mountoufaris et al., 2018) on the surfaces of individual neurons. What this code does during neural development is still being elaborated. In vivo, one of the most well-characterized functions for Pcdhs is in repulsion, both in the self-avoidance of same-cell dendrites and in repulsion or organization of certain axon tracts (Hasegawa et al., 2008, 2012, 2016; Katori et al., 2009, 2017; Prasad and Weiner, 2011; Chen et al., 2017; Lu et al., 2018). These are developmental processes that occur throughout embryonic and early postnatal development. However, Pcdhs are also highly expressed at later stages of development where synaptogenesis is prominent (Frank et al., 2005). The role of these molecules in synaptogenesis is currently unknown but some synaptic defects have been observed in Pcdh-*γ* null animals that differed depending on cell type (Weiner et al., 2005; Suo et al., 2012; Molumby et al., 2017). Here we find that hippocampal synapses with Pcdh-*γ* expression are significantly larger and have more perforations suggesting that these synapses may be more mature and/or stable.

Prior to synaptogenesis, dendrites elaborate an arbor that must occupy a distinct territory in the neuropil. In some neurons with planar dendritic trees, such as starburst amacrine cells in the retina or cerebellar Purkinje cells, same-cell dendrites rarely cross but can overlap with dendrites from neighboring neurons. It was shown in Pcdh-*γ* deficient mice that the dendrites from these neurons exhibited defective spatial distribution and abnormal clumping of the dendrites (Lefebvre et al., 2012; Ing-Esteves et al., 2018). In early stage hippocampal cultures, which are also planar, Pcdh-*γ*s align precisely at what have been termed “dendritic bridges” that could be morphological correlates of self-avoidance (Fernández-Monreal et al., 2009; Phillips et al., 2017). For avoidance to happen, it is almost certain that Pcdhs first make an adhesive linkage at contacting same-cell dendrites followed by detachment, cleavage or endocytosis of the adhesive complex in order for the contacting membranes to separate and dendrites to adopt different trajectories. These same-cell contacts are most likely to have a perfectly matching repertoire of Pcdhs. In contrast, Pcdh adhesive complexes at heterotypic junctions such as synapses or perisynaptic astrocyte contacts might have an “imperfect” match of Pcdhs that may behave differently than at same-cell contacts. Thus, our results, combined with others, suggests the possibility that Pcdhs may play different roles depending on context, with an avoidance role for dendrites of certain neurons and adhesive role in other instances.

During synaptic development, synapses make an initial adhesive contact through an expanding array of adhesion molecules (Kim, 2019). During postnatal life, robust synaptogenesis occurs but many presumptively functional synapses are also pruned. One possibility, based on their role in repulsion of same-cell dendrites, is that Pcdh adhesion participates in the synaptic pruning process. It would then be likely that synapses that exhibit Pcdh expression might exhibit a less mature phenotype than those that lack Pcdhs. The other possibility suggested by the present study is that Pcdh adhesion strengthens and stabilizes synaptic connections. It has been shown in vivo that the most persistent and stable spines have the widest diameter and a mushroom shape while transient spines are narrower (Berry and Nedivi, 2017). In the present study, synapses labeled with anti-Pcdh-*γ* were significantly wider and more likely to be perforated, both in vivo and in culture, and spine morphology was more likely to be of the mushroom type in culture.

### How might the endocytic trafficking model fit into the stabilization of synapses/spines?

Synapse stabilization is more consistent with an adhesive-like, rather than repulsive, role between pre- and postsynaptic neurons. Many of the synapses labeled in the present study were labeled intracellularly in addition to at the cleft. A model for Pcdh-mediated cell repulsion has been proposed in which an adhesive match between Pcdhs on adjacent cells induces rapid endocytosis which could remove other adhesive complexes from the cell surface and cause cell detachment (Phillips et al., 2017). How such a mechanism could be associated with synapse stabilization is not clear. Our results show that Pcdh-*γ*s are associated with more mature synapses but whether this involves a structural adhesive function is currently unknown. It is possible that endocytosis of Pcdhs after adhesive contact triggers a signaling mechanism that contributes to synapse stabilization. Overall our results do indicate that Pcdh-*γ*s are likely to be part of a synaptic maturation process with timely implications given the emerging role of Pcdh misregulation in neurodevelopmental disorders (El Hajj et al., 2017).

## Acknowledgements

Supported by R03 MH110980 to GRP.

## Notes

### Competing Interest Statement

The authors have declared no competing interest.

